# Cell cycle plasticity underlies fractional resistance to palbociclib in ER+/HER2- breast tumor cells

**DOI:** 10.1101/2023.05.22.541831

**Authors:** Tarek M. Zikry, Samuel C. Wolff, Jolene S. Ranek, Harris Davis, Ander Naugle, Austin A. Whitman, Michael R. Kosorok, Philip M. Spanheimer, Jeremy E. Purvis

## Abstract

The CDK4/6 inhibitor palbociclib blocks cell cycle progression in ER+/HER2- breast tumor cells. Although these drugs have significantly improved patient outcomes in metastatic breast cancers, a small percentage of tumor cells continues to divide in the presence of palbociclib—a phenomenon we refer to as fractional resistance. It is critical to understand the cellular mechanisms underlying fractional resistance because the precise percentage of resistant cells in patient tissue is a strong predictor of clinical outcome. Here, we hypothesize that fractional resistance arises from cell-to-cell differences in core cell cycle regulators that allow a subset of cells to escape CDK4/6 inhibitor therapy. We used multiplex, single-cell imaging to identify fractionally resistant tumor cells both in a cell culture model of ER+/HER2- breast cancer as well as live primary tumor cells resected from a patient. We found that tumor cells capable of proliferating in the presence of palbociclib showed both expected (e.g., CDK2, E2F1) and unexpected (e.g., Cdt1, p21, cyclin B1) shifts in core cell cycle regulators. Notably, resistant cells in both tumor models showed premature enrichment of the G1 regulators E2F1 and CDK2 protein and, unexpectedly, the G2/M regulator cyclin B1 just before cell cycle entry, suggesting that resistant cells may use noncanonical mechanisms to overcome CDK4/6 inhibition. Using computational data integration and trajectory inference approaches, we show how plasticity in cell cycle regulators gives rise to alternate cell cycle “paths” that allow individual ER+/HER2- tumor cells to escape palbociclib treatment. Understanding drivers of cell cycle plasticity, and how to eliminate resistant cell cycle paths, could lead to improved cancer therapies targeting fractionally resistant cells to improve patient outcomes.

## INTRODUCTION

Estrogen receptor-positive, human epidermal growth factor 2 receptor-negative (ER+/HER2-) metastatic breast cancers shows altered cell cycle behaviors that contribute to progression of the disease (1–3). Most ER+/HER2- tumors show elevated expression of the estrogen receptor (ER, *ESR1*) and its transcriptional target, cyclin D1 (*CCND1*) (4–6). Estrogen signaling upregulates expression of cyclin D1, which works together with other cyclins (e.g., cyclin E) to activate cyclin dependent kinases (CDKs) (7). Cyclin D1 forms complexes with CDK4 and CDK6 (8, 9), whereas cyclin E forms complexes with CDK2. Active cyclin-CDK complexes phosphorylate the retinoblastoma protein (RB) to its phosphorylated form (pRB) (10, 11). RB phosphorylation relieves repression of a large set of target genes controlled by the E2F family of transcription factors (e.g., E2F1) (12–14). Expression of E2F-regulated genes produces additional positive feedback mechanisms to initiate S phase (13, 15). In general, ER+/HER2- breast tumors show enhanced signaling through cyclin-CDK signaling pathways that converge on phosphorylation of RB to initiate cell cycle entry and tumor cell proliferation (**Figure 1A**).

**Figure 1.**
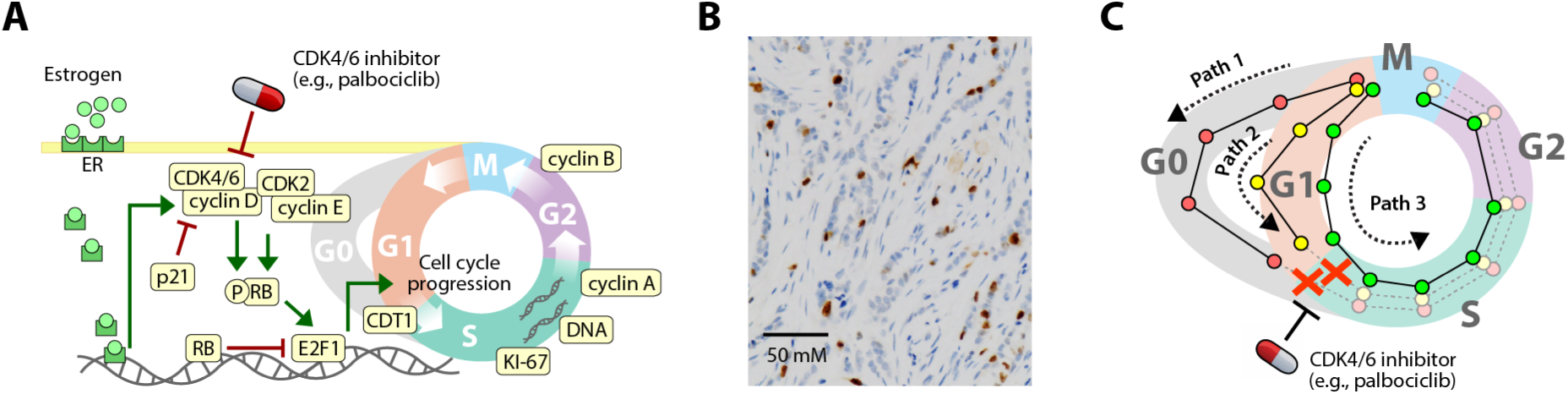
Cell cycle regulation and fractional resistance to CDK4/6 inhibitors ER+/HER2- breast tumor cells. **A.** Core cell cycle signaling network and points of drug activity in ER+/HER2- breast tumors. Cell cycle protein regulators shown in beige were measured in single tumor cells. **B.** KI-67 staining *(brown)* from an ER+/HER2- breast tumor with 14% KI-67+ cells. **C.** Hypothetical model in which plasticity in cell cycle progression among individual cells could create different molecular “paths” with distinct sensitivities to CDK4/6 inhibitors. Red, yellow, and green paths represent distinct molecular trajectories for a single cell. In the proposed model of fractional resistance, CDK4/6 inhibitors do not block all paths, allowing some cells to complete the cell cycle in the presence of the drug.

For over twenty years, the mainline treatments for ER+/HER2- breast tumors have been endocrine therapies such as tamoxifen, aromatase inhibitors, and fulvestrant. These “anti-estrogen” agents partially block or antagonize estrogen receptor signaling to reduce cyclin D1 expression. More recently, several potent CDK4/6 inhibitors such as palbociclib, abemaciclib, or ribociclib, are being given in combination with endocrine therapy to improve patient outcomes. These drugs have had a profound clinical impact. Data from multiple clinical trials shows that combined anti-estrogen and CDK4/6 inhibitor therapy nearly doubles progression-free survival, and many patients respond to CDK4/6 inhibitor therapy for several years (6, 16–18). However, many patients are initially resistant to the CDK4/6 inhibitors or acquire resistance within the first few months of treatment. Despite extensive investigation, the mechanisms of resistance to CDK4/6 inhibitors in ER+/HER2- breast cancers remain unclear (19, 20).

An important clue in understanding resistance to CDK4/6 inhibitors and endocrine therapy comes from Ki-67 staining, a common clinical diagnostic for breast tumors. Ki-67 is a nuclear protein that is expressed in actively proliferating cells but is absent in arrested cells (21). Ki-67 staining in paraffin-embedded tissues nearly always shows a subpopulation of proliferating cells (21, 22) (**Figure 1B**). This percentage of proliferating cells is strongly predictive of clinical outcomes. After receiving treatment, for example, patients showing a low percentage (0-2%) of proliferating cells are considered “responsive” to therapy, whereas a higher percentage (3-15%) of Ki-67-positive cells predicts a poor response for ER+/HER2- patients (23). Recent work by Gaglia *et al.* shows that Ki-67 likely underestimates the number of proliferative cells in tumor tissues (24). These observations strongly suggest that a subpopulation of tumor cells continues to divide, even in “responsive” ER+/HER2- patients. We refer to this phenomenon in which a subpopulation of tumor cells continues to divide in the presence of a cell cycle-targeting drug as fractional resistance. Fractional resistance is conceptually similar to fractional killing, a term coined to describe the observation that each round of chemotherapy does not kill 100% of tumor cells (25). Previous work (26–34) showed that fractional killing is a consequence of non-genetic, cell-to-cell heterogeneity—that is, differences in the molecular makeup of individual cells that do arise from genetic mutations. Additional studies indicate that cell-to-cell heterogeneity plays a role in resistance in melanoma (28, 35, 36).

One source of cell-to-cell heterogeneity is cell cycle plasticity—differences in the cell cycle behavior driven by different combinations of cell cycle regulators. A recent meta-analysis of cancer cell lines by Kumarasamy and colleagues found that sensitivity to CDK4/6 inhibitors was associated with activation of RB, inhibition of CDK2 activity and, in some cases, depletion of CDK4 and CDK6 (37). These results demonstrate how cell cycle plasticity across different cell lines leads to drug resistance, but it does not indicate what cell cycle plasticity may exist within a single type of tumor cell, potentially explaining its drug resistance. A growing body of evidence (37–43), including our own work (44–47), has shown that differences in cell cycle behavior occurs at the level of individual cells. These differences include changes in the timing of key cell cycle events such as a shortened G1 duration (42, 44, 46, 48) as well as an altered ordering of events at the G1/S transition (49, 50). The cell cycle can also vary at the single-cell level in its pattern of cyclin expression (51), CDK activity (15, 38), and other molecular states (37, 43). These studies strongly suggest that individual cells can take distinct trajectories, or “paths”, through the cell cycle that are defined by a unique combination of molecular states over time (52, 53) (**Figure 1C**).

Here, we show that ER+/HER2- tumor cells show subtle, cell-to-cell differences in core cell cycle regulators that allows a subset of tumor cells to escape CDK4/6 inhibitor therapy. We used multiplex, single-cell imaging to build a proteomic profile for individual tumor cells treated with palbociclib and identified fractionally resistant tumor cells both in a cell culture model of ER+/HER2- breast cancer as well as from live primary tumor cells resected from a patient. We found that resistant tumor cells harbored specific combinations of enriched and depleted cell cycle regulators including cyclin B1, Cdt1, CDK2, and p21, some of which were common to both tumor models. Using computational data integration and trajectory inference approaches to visualize resistant cells, our work shows how non-genetic plasticity in cell cycle regulators—at the single-cell level—creates alternate cell cycle paths that allow individual ER+/HER2- tumor cells to escape palbociclib treatment.

## RESULTS

We first investigated fractional resistance in the T47D cell line, a well-established model of ER+/HER2- breast cancer (54, 55). Cells were allowed to proliferate freely or treated with either low (10 nM) or moderate (100 nM) doses of palbociclib for 24 hours. We then profiled protein expression in 103,862 cells using indirect iterative immunofluorescence imaging, or “4i” (56) (**Figure 2A**). For each cell, we quantified the abundance of 14 cell cycle regulators: pRB, RB, Ki-67, CDK2, CDK4, cyclin D1, cyclin E, Cdt1, E2F1, cyclin A, cyclin B1, p21, and integrated DNA (**Table S1**). These core regulators cover a broad range of molecular mechanisms occurring throughout the cell cycle, including growth signaling (cyclins D and E, CDKs 2 and 4), the G1/S transition (RB, pRB, E2F1), DNA replication (Cdt1, Ki-67, DNA), and progression through G2 and M phases (cyclins A and B). Given that a freely dividing population of tumor cells is not synchronized to any particular cell cycle phase, 4i captures a diverse range of single-cell states across all phases of the entire cell cycle. To facilitate a direct comparison of cell cycle states across samples and treatment conditions, we performed principled downsampling of the data using kernel herding sketching (57). This approach identifies a limited subset of representative cells that preserves the original distribution of cell states. We selected an equal number of cells (*n* = 2000) from each treatment condition (untreated, 10 nM, 100 nM), resulting in a final downsampled dataset of 6,000 T47D cells (see **MATERIALS AND METHODS**).

**Figure 2.**
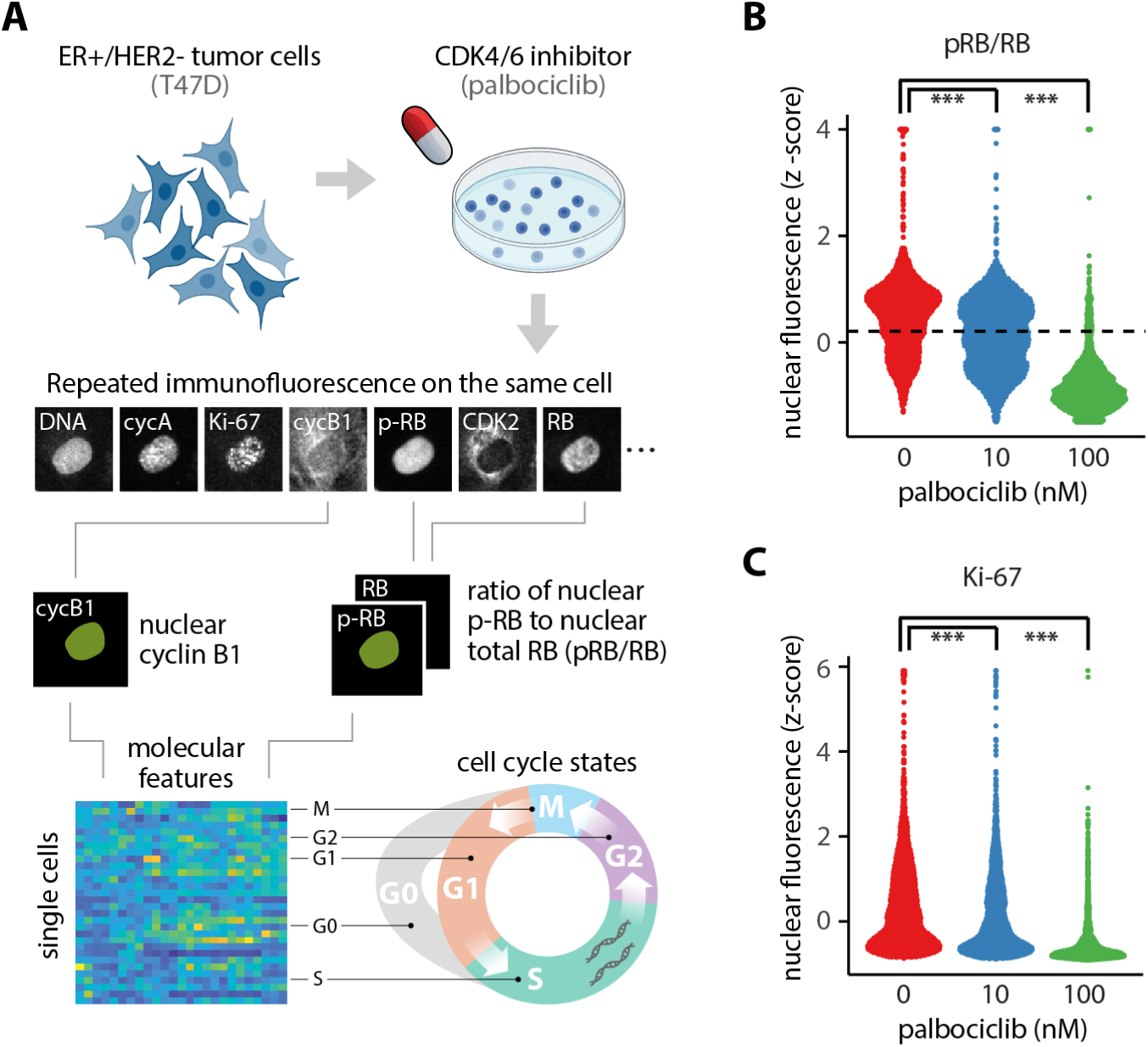
Single-cell proteomic profiling reveals fractional resistance to palbociclib in ER+/HER2- breast tumor cells. **A.** Workflow for 4i profiling of breast tumor cells. Asynchronous T47D cells were treated with increasing concentrations of palbociclib for 24 hours. Cells were fixed and subjected to iterative indirect immunofluorescence imaging (4i) to quantify nuclear levels of pRB, RB, Ki-67, CDK2, CDK4, cyclin D1, cyclin E, Cdtl, E2Fl, cyclin A, cyclin Bl, p21, and DNA in each cell. The resulting data structure is a matrix containing 103,862 cells and imaging features representing the 14 cell cycle regulators. Because the cells are not synchronized, 4i capturesa highly granular representation of proliferating and arrested cell cycle states. **B.** Distribution of pRB/RB at 0, 10, or 100 nM palbociclib. The dotted line marks the statistically determined threshold for demarcating cells as proliferating despite palbociclib treatment. Cells above the dotted line were used for characterizing proliferating cells. **C.** Distribution of nuclear Ki-67 levels at 0, 10, or 100 nM palbociclib. *** indicates a P-value < 0.001 using a two-sided Kolmogorov-Smirnov test between untreated and treated cells.

To quantify the fraction of proliferating cells under each condition, we focused on the ratio of phosphorylated to total RB protein, or pRB/RB. When quantified in individual cells, pRB/RB often shows a bimodal distribution: low pRB/RB expression corresponds to a hypo-phosphorylated RB state and is characteristic of arrested cells. High pRB/RB expression represents the hyper-phosphorylated form of the RB protein and comprises actively proliferating cells (58). As expected, palbociclib reduced the fraction of proliferating cells in a dose-dependent manner (**Figure 2B**). Interestingly, however, we observed a small subset of cells that maintained high pRB/RB levels at both 10 nM (48.4%) and 100 nM (5.9%) palbociclib, indicating that some cells could evade drug treatment. Similarly, we observed a reduction in Ki-67 expression under increasing palbociclib concentration, but a fraction of cells maintained high Ki-67 expression in the presence of 10 nM and 100 nM palbociclib (**Figure 2C**). We observed a similar level of fractional resistance in a biological replicate of T47D cells (**Figure S1A-B**). To confirm the cells were truly capable of proliferating in the presence of palbociclib, and not merely finishing the previous cell cycle within the 24-hour treatment time frame, we repeated the experiment in long-term culture, exposing cells to one week of continuous palbociclib treatment. Again, tumor cells showed fractional resistance as indicated by the continual presence of proliferating cells (**Figure S1C**). Taken together, these results reveal that a small subset of ER+/HER2- breast tumor cells continue to proliferate in the presence of palbociclib, implying that cell-to-cell variation may account for fractional resistance.

We next asked what intracellular features may be enriched in proliferating cells in the presence of palbociclib. We focused deliberately on differences in protein abundance—rather than epigenetic or transcriptomic differences—because palbociclib acts directly on the CDK4/6 proteins (59), and because the core cell cycle regulators are largely regulated through protein modification and degradation (60, 61). To identify only the proliferating cells, and exclude G0 cells, we set a threshold level of pRB/RB above which cells were confidently expected to be in the hyperphosphorylated state and therefore actively proliferating in either G1, S, G2, or M phases (38, 62, 63). We set a conservative cutoff (see **MATERIALS AND METHODS**) to capture this second peak of pRB/RB expression (dotted line in **Figure 2B**). We then compared the single-cell profiles between these proliferating untreated and palbociclib-treated cells to identify differences in cell cycle regulators that may be responsible for fractional resistance. Shifts in the distributions of individual cell cycle proteins (**Figure 3A**) were quantified using a two-sample *t-*-test between untreated cells and either the 10 nM or the 100 nM treatment condition, producing 95% confidence intervals (CIs) for any observable differences in single-cell protein expression (**Figure 3B**).

**Figure 3.**
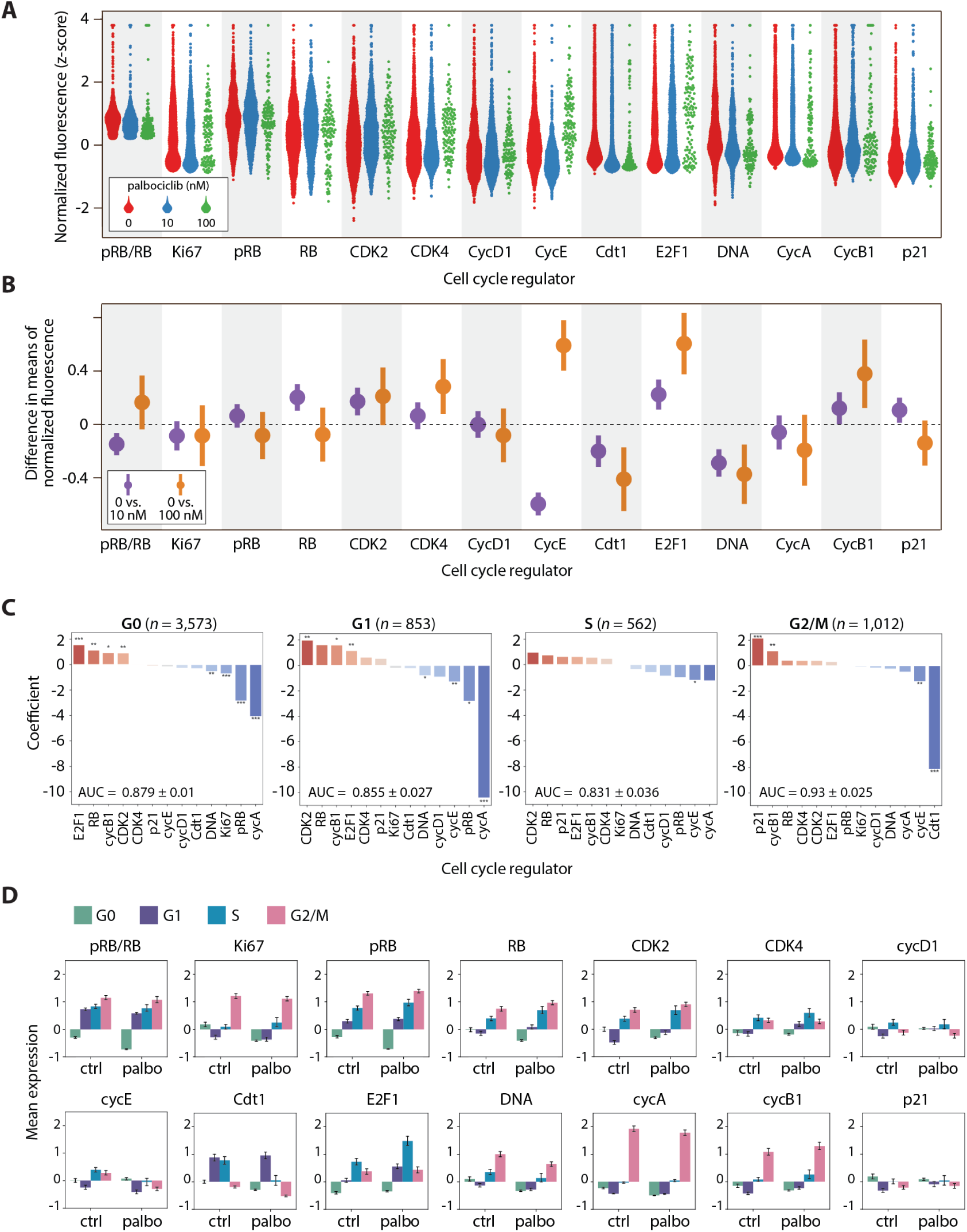
Shifts in expression of cell cycle proteins among fractionally resistant tumor cells. **A.** Beginning with a downsampled dataset containing 2,000 cells from each condition (see **MATERIALS AND METHODS),** we identified the proliferating cells using the pRB/RB threshold defined in Figure 2B. Proliferating single-cell distributions of cell cycle regulators at O nM, 1O nM, or 100 nM palbociclib, showing an expected reduction in the number of cells at higher drug concentrations. **B.** 95% confidence intervals (Cl) of proliferating cells for the differences in mean expression (normalized z-scores) between either untreated cells and 1O nM palbociclib *(purple);* or untreated and 100 nM palbociclib *(orange).* Confidence intervals overlapping with the dashed line at O indicate a lack of statistical significance. Confidence intervals are wider when comparing O vs. 100 nM due to lower sample sizes at the highest dose of palbociclib. **C.** Logistic regression on all 6,000 cells predicting the odds that a given cell is either untreated (O nM) or treated (1O nM or 100 nM) based on expression of its cell cycle regulators. Cell-to-cell increases in regulators shown in red (e.g., CDK4), or decreases in regulators shown in blue (e.g., CDK2), increase the odds of association with treated (10 nM or 100 nM) versus untreated cells. A separate regression was performed for each phase (GO, Gl, S, G2/M), where the last three are considered proliferating (high pRB/RB). This analysis was performed on all cells, including non-proliferating (GO) cells. Significance:*, *P* < 0.05; **, *P* < 0.01, ***, *P* < 0.001. **D.** Expression levels of cell cycle features, stratified by cell cycle phase for all 6,000 cells. Bar height is the mean expression for untreated (ctrl) or palbociclib-treated (1O nM and 100 nM) cells. Error bars represent confidence intervals.

Under palbociclib treatment, many cell cycle regulators (e.g., pRB, Ki-67, cyclin A) showed similar distributions of expression compared to untreated cells. However, some regulators showed significant shifts in protein expression under 10 nM and/or 100 nM palbociclib treatment. For example, proliferating T47D cells showed elevated CDK2 levels under 10nM and 100 nM palbociclib treatment (0 vs. 10 nM 95% CI (0.09, 0.26); 0 vs. 100 nM (0.014, 0.41)) and reduced expression of Cdt1 (0 vs. 10 nM (−0.30, −0.10,); 0 vs. 100 nM (−0.63, −0.19) at both drug doses. Elevated expression of CDK2 activity is consistent with previous studies showing that ER+/HER2- tumors cells often become resistant to CDK4/6 inhibitors via increases in cyclin E/CDK2 activity (5, 16). However, we note that these studies are typically focused on genetic changes in cyclin E/CDK2 activity (e.g., mutations or copy number variation) and not cell-to-cell variability in protein expression. On the other hand, reduced Cdt1 expression under palbociclib treatment has, to our knowledge, not been previously described. Cdt1 encodes for a subunit of the pre-replication complex necessary for DNA replication. Entry into S phase with sub-normal Cdt1 levels could potentially lead to replication stress (64). Finally, we noted that proliferating T47D cells showed a significant depletion of DNA content, likely reflecting a relative enrichment of G1-arrested cells with elevated pRB/RB levels.

We next asked how expression of cell cycle regulators shifted as cells progressed through individual cell cycle phases. To do this, we performed unsupervised clustering of the expression levels of DNA, cyclin A, and cyclin B1 to assign each cell to a specific cell cycle phase: G0, G1, S, or G2/M (**MATERIALS AND METHODS**). We then trained a logistic regression model within each phase separately on the proteomic expression profiles from all 6,000 cells to classify whether a cell belonged to either the control (untreated) or treatment (10 nM and 100 nM palbociclib) group (**Figure 3C**). When paired with a raw comparison of changes in expression (**Figure 3D**), this analysis reveals how differences in expression of cell cycle regulators—either positive changes (red bars in **Figure 3C**) or negative changes (blue bars in **Figure 3C**)—are associated with cell cycle progression under palbociclib treatment. The results were largely consistent with the previous *t*-test analysis of proliferating cells. For example, we found that elevated CDK2 expression was a significant, positive predictor of palbociclib-treated cells in both G0 and G1 phases. However, this analysis also revealed several new trends. For example, increases in three other core cell cycle regulators—E2F1, RB, and cyclin B1—were significantly associated with palbociclib-treated cells in G0 and/or G1 phases. E2F1 expression was among the strongest predictors of treatment in both G0 and G1 cells, providing further clarity for the overall E2F1 enrichment we observe under treatment in **Figure 3B**. These results suggest that individual tumor cells with prematurely elevated E2F1 and CDK2 protein levels may be more likely to transition from G0 to G1 in the presence of palbociclib.

More surprisingly, we found that enrichment of cyclin B1 was positively associated with palbociclib-treated cells in G0, G1, and G2/M phases. Cyclin B1 is a G2 cyclin that is required for entry into M phase (65). The untimely enrichment of cyclin B1 among treated cells in G0 and G1 phases suggests that elevated cyclin B1 levels may facilitate resistance to palbociclib through noncanonical mechanisms of cell cycle progression. Enrichment of cyclin B1 in G0 and G1 phases was confirmed after performing the same analysis on a biological replicate of T47D cells (**Figure S2C**). Overall, we found that fewer cell cycle regulators reached statistical significance in S or G2/M phases than in G0 or G1 phases, likely because of the smaller sample sizes for these subpopulations of cells, especially under palbociclib treatment where most cells were arrested in G0. Nevertheless, several overall trends were consistent across all phases; specifically, these phase-specific analyses show that palbociclib either promotes, or selects for, proliferation of tumor cells with altered protein expression profiles characterized by increases in E2F1, CDK2, and cyclin B1, and decreases in Cdt1.

Although T47D is a well-established model of ER+/HER2- breast cancer, it may not reflect the physiology of primary tumor cells from an actual breast cancer patient. Thus, we developed an experimental strategy for studying fractional resistance in surgically resected primary human tumors (**Figure S3**, **MATERIALS AND METHODS**). Briefly, we obtained a specimen from a primary ER+/HER2- invasive lobular carcinoma which was delivered immediately from the operating room to the laboratory. After dissociation, cells were plated, treated with palbociclib or control media for 24 hours, and subjected to 4i profiling. Because tissue samples contain complex mixtures of cell types, we used expression of the epithelial marker pan-cytokeratin (PanCK), as well as the estrogen and progesterone receptors (ER/PR) to computationally separate the tumor cells for downstream analysis. Out of a total population of 100,191 cells, we identified a subpopulation of 14,789 cells showing high expression of ER, PR, and PanCK (**Figure S4**). As above, we performed principled downsampling to a size of 6,000 cells (2,000 cells per condition) to match the T47D analysis.

We first looked for shifts in protein expression between untreated and palbociclib-treated primary tumor cells. As before, we focused initially only on the proliferating subpopulations (i.e., G1, S, and G2/M cells) using a conservative cutoff for pRB/RB levels to identify fractionally resistant tumor cells (**Figure S5**). Although we did not observe consistent changes in Cdt1 or CDK2 expression, primary tumor cells shared many of the same shifts in expression as T47D cells. For example, we observed a significant depletion of DNA content (0 vs. 100 nM 95% CI (−0.89, −0.21)) and additionally, Ki-67 expression (0 vs. 100 nM 95% CI (−0.95, −0.12)) (**Figure 4B**). As with T47D, the depletion of Ki-67 and DNA was likely due to the accumulation of G1 cells under treatment, which would be predicted to have lower DNA and Ki-67 content (66). We also observed a significant upregulation of E2F1 protein levels, as previously observed in T47D. A distinct finding in primary tumor cells was the consistent and significant depletion of cyclin A (95% CI (−1.12, −0.54)) and p21 (95% CI (−0.80, −0.28)) under palbociclib treatment. The downward shift in cyclin A could be due to a depletion of S and G2/M cells relative to G1 cells among proliferating treated cells.

**Figure 4.**
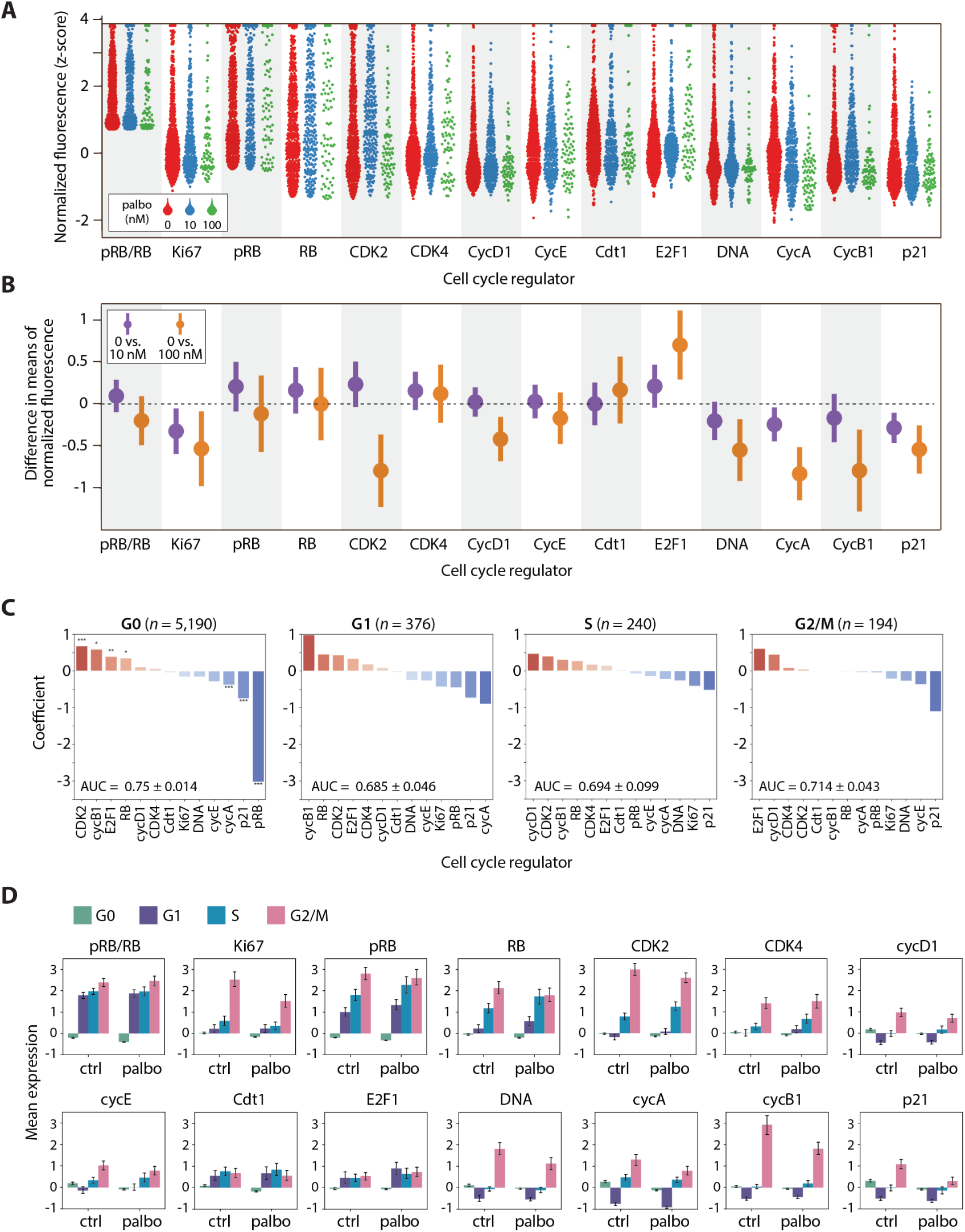
Palbociclib reveals shifts in expression of cell cycle proteins in primary tumor cells. **A.** As with T47D, we identified proliferating primary tumor cells using the pRB/RB threshold defined in **Figure S5**. Proliferating single-cell distributions of cell cycle regulators at O nM, 1O nM, or 100 nM palbociclib, showing an expected reduction in the number of cells at higher palbociclib concentrations. **B.** 95% confidence intervals (Cl) of proliferating cells for the differences in mean expression (normalized z-scores) between either untreated cells and 1O nM palbociclib *(purple);* or untreated and 100 nM palbociclib *(orange).* Confidence intervals overlapping with the dashed line at O indicate a lack of statistical significance. Confidence intervals are wider when comparing O vs. 100 nM due to lower sample sizes at the highest dose of palbociclib. **C.** Logistic regression on all 6,000 cells predicting the odds that a given cell is either untreated (O nM) or treated (1O nM or 100 nM) based on expression of its cell cycle regulators. Cell-to-cell increases in regulators shown in red (e.g., CDK2), or decreases in regulators shown in blue (e.g., p21), increase the odds of association with treated (10 nM or 100 nM) versus untreated cells. A separate regression was performed for each phase (GO, G1, S, G2/M), where the last three are considered proliferating (i.e., high pRB/RB).This analysis was performed on all cells, including non-proliferating (GO) cells. Significance:*, *P* < 0.05; **, *P* < 0.01, ***, *P* < 0.001. **D.** Expression levels of cell cycle features, stratified by cell cycle phase for all 6,000 cells. Bar height is the mean expression for untreated (ctrl) or palbociclib-treated (10 nM and 100 nM) cells. Error bars represent confidence intervals.

Although CDK2 showed an inconsistent pattern of expression characterized by apparent enrichment at 10 nM and depletion at 100 nM palbociclib treatment, this discrepancy was clarified by the logistic regression model, which identified elevated CDK2 expression as a top-ranking predictor of palbociclib treatment across all cell cycle phases (**Figure 4C**). By examining the average change in expression for each cell cycle regulator across the cell cycle phases (**Figure 4D**), we observed a gradual accumulation of CDK2 as cells progressed from G1, through S phase, peaking in G2/M, consistent with the protein’s known dynamical pattern of expression (67, 68). Thus, accumulation of CDK2 protein in palbociclib-treated tumor cells suggests a potential axis of nongenetic resistance mediated by enhanced cyclin E/CDK2 activity signaling.

A more striking consistency with T47D cells was that the same four cell cycle regulators— CDK2, cyclin B1, E2F1, and RB—were again the highest-ranking and most significant positive factors associated with G0 arrest in primary tumors cells. This result points to a potentially common mechanism of non-genetic drug resistance. In addition, we also noted that increased cyclin B1 expression in G0 cells was positively associated with palbociclib treatment (**Figure 4C**), consistent with results in T47D. At first, this result seemed to contradict the results shown in **Figure 4B**, where cyclin B1 is apparently decreasing under palbociclib treatment. However, a closer examination of average cyclin B1 expression in **Figure 4D** shows that the decrease is driven primarily by G2/M cells, the least populous phase of tumor cell (see pie charts in **Figure 5C**) and the remaining phases are either increasing in expression of cyclin B1 under treatment, or staying at the same level. Furthermore, as there are fewer proliferating cells under 100 nM of palbociclib than at 10 nM, the apparent decrease under the higher dose may be lost by grouping treatment conditions.

**Figure 5.**
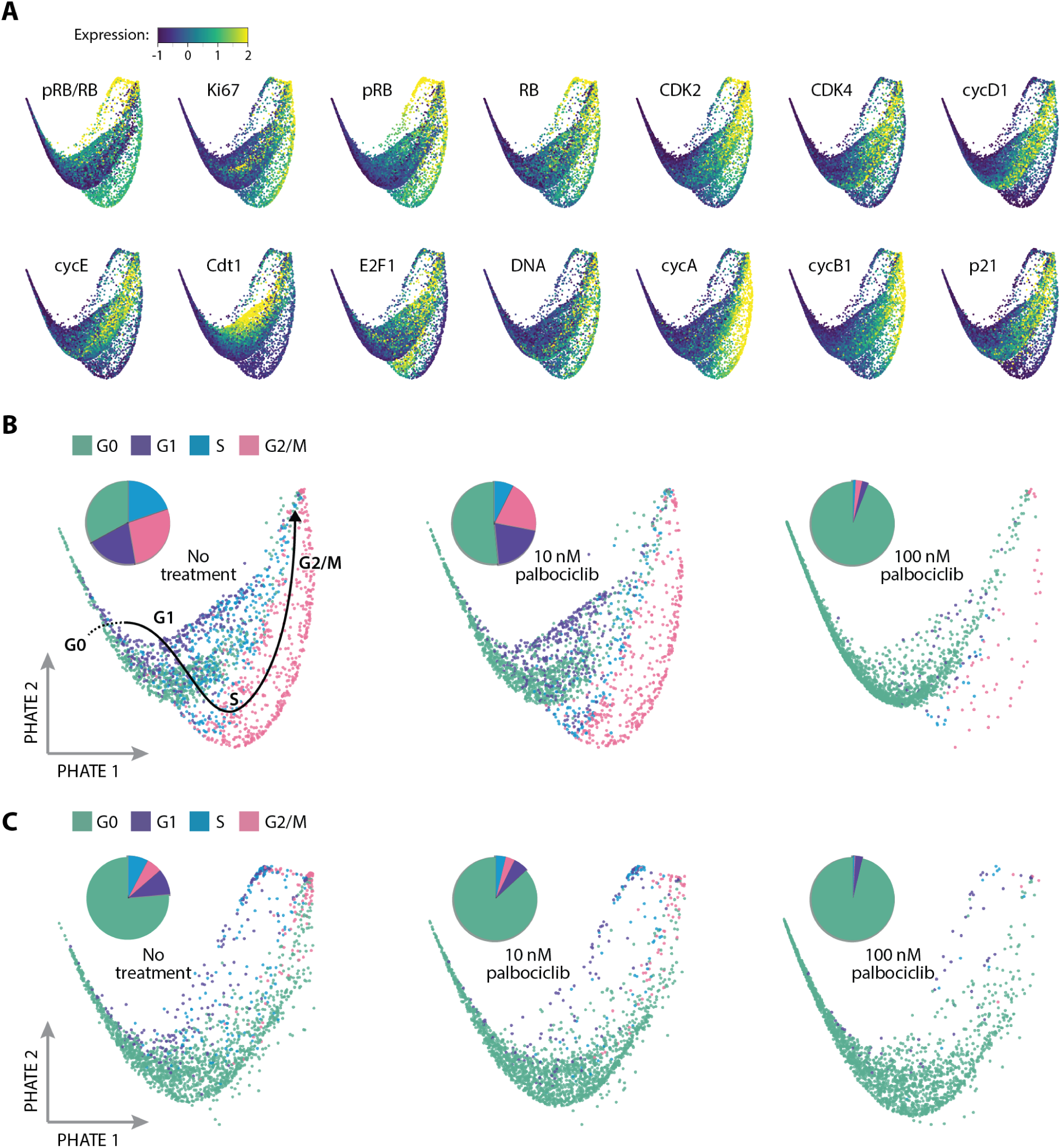
Visualization of single cell states in ER+/HER2- breast tumors under palbociclib treatment. T47D and primary tumor samples were independently downsampled to yield a sample size of 6,000 cells each (2,000 cells from each of the three palbociclib treatment conditions). The datasets were then integrated into a joint latent space of 12,000 cells using TRANSACT. We then applied the nonlinear dimensionality method PHATE to produce a 2-dimensional visualization of cell cycle under each condition. Each dot is an individual cell. **A.** Expression levels of each cell cycle regulator overlaid onto the PHATE embedding for both tumor models. **B.** Visualization of cell cycle states for T47D cells, separated by treatment condition. Pie charts indicate the proportion of cells in each cell cycle phase. A hand-drawn estimate of cell cycle progression is shown on the first image based on the progression of overlaid features in panel **A. C.** Visualization of cell cycle states for primary tumor cells, separated by treatment condition.

Interestingly, the cell cycle inhibitor, p21, which serves to block phosphorylation of RB by inhibiting cyclin-CDK complexes, was significantly downshifted under both 10 nM and 100 nM palbociclib treatment (**Figure 4B**, 0 vs. 10 nM 95% CI (−0.44, −0.13), 0 vs. 100 nM (−0.79, −0.28)). The logistic regression model also identified p21 as a strongly negative predictor of palbociclib treatment across all cell cycle phases (**Figure 4C**), and p21 showed a consistent pattern of reduced expression across all phases (**Figure 4D**). This result was not observed for T47D cells. The palbociclib-induced reduction in p21 levels could potentially relax cell cycle arrest and allow cells to enter G1 with modest cyclin/CDK activity. This observation is consistent with enrichment of E2F1 under palbociclib treatment, which was significantly upshifted in primary human tumor cells. Despite these discrepancies among tumor models, consistent enrichment of CDK2, E2F1 and cyclin B1 in G0 and G1 phases, both in T47D and primary tumor cells, suggests a potentially common mechanism of fractional resistance. The accumulation of such factors potentially allows a subpopulation of cells to prematurely enter G1 with elevated pRB/RB levels. These findings indicate that the tumor cell cycle is inherently plastic; individual cells can take different molecular paths through the cell cycle, some of which are resistant to CDK4/6 inhibitors.

We next sought to visualize resistant cell cycle paths taken by individual tumor cells. To this end, we first identified a shared latent space of T47D and primary tumor samples by performing data integration with Tumor Response Assessment by Nonlinear Subspace Alignment of Cell lines and Tumors (TRANSACT) (69) (see **MATERIALS AND METHODS**). We then applied Potential of Heat-diffusion for Affinity-based Trajectory Embedding (PHATE) (70) on the joint latent space to generate a low-dimensional projection of the combined tumor cell cycle (52, 53). **Figure 5** shows the resulting saddle-shaped structure that captured the progression of four cell cycle phases and revealed significant variability in single cell states. Each dot in the structure represents an individual cell, and dots nearer to each other have similar expression profiles of cell cycle regulators. By overlaying expression levels for individual cell cycle regulators (**Figure 5A**), we observed well-established cell cycle events, including elevated pRB and Ki-67 in proliferating cells; a peak in Cdt1 expression in late G1; accumulation of E2F1 in S phase; and sequential expression of cyclin A and cyclin B1. These temporal trends in core cell cycle regulators, which emerged without providing any input to the model, allowed us to estimate the general path of cells from G0 to G1, S, and G2/M phases (back curve in **Figure 5B**). As expected, we found that palbociclib gradually depleted the number of cells in proliferative phases for both T47D (**Figure 5B**) and primary tumor cells (**Figure 5C**), potentially altering the path of proliferation through the cell cycle.

To obtain a more objective and quantitative calculation of the various paths cells take through the cell cycle under palbociclib treatment, we performed trajectory analysis using Slingshot (71) through the low-dimensional PHATE embedding. Without specifying the correct order of phases, this analysis determined the proper cell cycle ordering from G0, to G1, to S, and to G2/M phase in most of the trajectories (see **MATERIALS AND METHODS**). To allow direct comparisons between each path, we aligned the separate treatment trajectories into one shared axis for each data source using TrAGEDy (72). **Figure 6** shows a visualization of the different temporal expression trends of cell cycle regulators under each experimental condition. In both T47D and the primary tumor cells (**Figure 6A, D**), we observed a clear inward movement of the trajectories—away from proliferating subpopulations—as palbociclib dose increased. Heatmaps in **Figures 6B** and **6E** illustrate the scaled expression of cell cycle regulators ordered along the aligned pseudotime axis. As an alternative visualization, **Figures 6C** and **6F** directly compare the aligned pseudotime traces for each cell cycle regulator across treatment conditions for T47D and primary tumor samples, respectively. To facilitate comparison among the drug doses, we defined a common transition point as the time at which 50% of the cells remained in G0 and 50% of the cells had progressed to a proliferative phase (vertical lines in panels **B**, **C**, **E**, and **F**).

**Figure 6.**
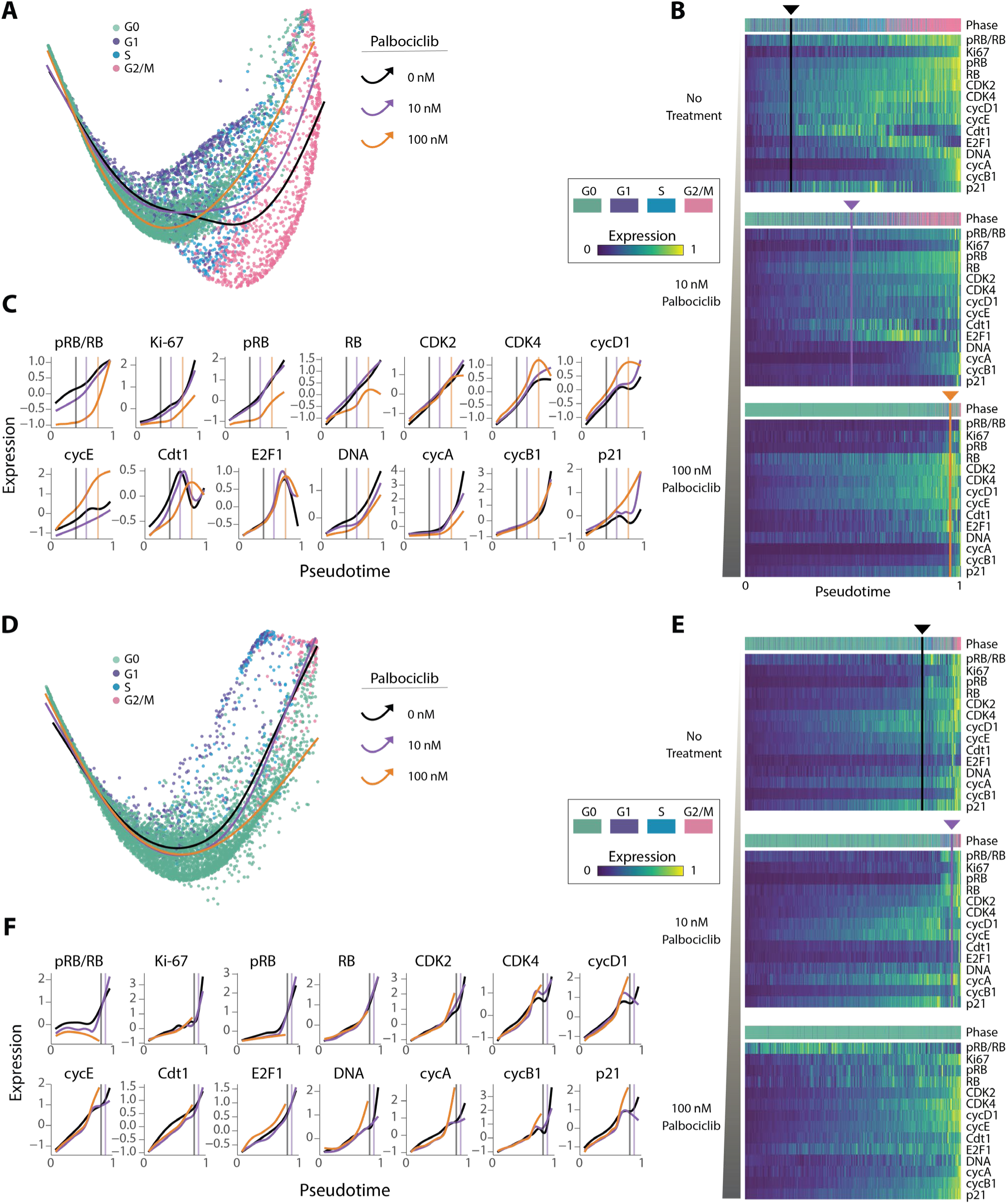
ER+/HER2-tumor cells take alternate cell cycle trajectories under palbociclib treatment. Trajectory inference was performed on cells from each treatment condition forT47D (**A-C**) and primary tumor (**D-F**) samples using Slingshot on the joint PHATE embedding. Each trajectory started in GO, progressed through the proliferative phases of the cell cycle, and ended in G2/M. A common pseudotime axis (**B, C, E, F**) was determined by aligning trajectories across treatment conditions using TrAGEDy (see **MATERIALS AND METHODS**). **A, D.** Trajectories for each treatment condition projected onto the joint two-dimensional PHATE embedding. **B, E.** Heatmaps showing expression of cell cycle regulators along the pseudotime trajectories in panel **A.** The color strip above the heatmaps represents the cell cycle phase annotations for each cell in the pseudotime ordering. Vertical lines indicate the time at which approximately half of the cells have transitioned from GO to a proliferative phase (G1, S, G2/M). No line was detectable for primary tumor cells under 100 nM palbociclib treatment because too few cells entered proliferative phases. **C, F.** Comparison of trajectories across treatment conditions for each cell cycle regulator. Vertical lines mark the same common transition point from GO to proliferation for each drug dose.

As expected, the temporal trends of increasing pRB/RB and Ki-67 levels, representing escape from cell cycle arrest, were delayed (i.e., right-shifted) under palbociclib treatment in a dose-dependent fashion (**Figures 6C** and **6F**). Despite these delays in cell cycle entry, however, many other cell cycle regulators nevertheless maintained synchronized trends. For example, in both tumor models, CDK2, E2F1, and cyclin B showed temporally synchronized increases in expression among all treatment groups. Importantly, however, in palbociclib-treated cells, these factors accumulated for longer times and to greater extents before the population reached the G0/G1 transition point, when approximately half of the cells were determined to begin proliferation. More strikingly, other cell cycle regulators showed a reversal of temporal trends. For example, T47D cells treated with 100 nM of palbociclib showed early increases in CDK4, cyclin D1, and cyclin E compared to 10 nM and untreated cells. Conversely, Cdt1 expression was both reduced and delayed upon drug treatment (**Figure 6A-C**). In contrast to T47D, in the primary ER+/HER2- tumor (**Figure 6D-F**), expression of p21 was reduced among treated cells. Taken together, this analysis supports the observation that accumulation of distinct cell cycle-promoting factors, including E2F1, CDK2, and the G2-associated cyclin B1, precedes—and potentially facilitates—escape from palbociclib-mediated arrest in T47D and primary tumor cells.

## DISCUSSION

In ER+/HER2- breast cancer, drugs that specifically inhibit cyclin-dependent kinases 4 and 6 (CDK4/6 inhibitors)—when given in combination with endocrine therapy—have dramatically improved oncologic outcomes and overall survival. Unfortunately, there is considerable heterogeneity in the clinical responses to CDK4/6 inhibitors, and most patients eventually develop drug resistance. The field has poured tremendous effort into genetic profiling studies with the hope of identifying molecular mechanisms that predict resistance to CDK4/6 inhibitors. Besides identifying a handful of genes associated with resistance, there is currently no biomarker in clinical use that can predict how an ER+/HER2- patient will respond to endocrine therapy and CDK4/6 inhibitors. This is not just a failure of precision medicine—it also reveals a serious gap in our understanding of the mechanisms that underlie drug resistance.

Here, we provide a new framework for how drug resistance may arise in ER+/HER2- breast tumors. We show that cell-to-cell differences in core cell cycle regulators allow a subset of tumor cells to escape CDK4/6 inhibitor therapy. We refer to this phenomenon as fractional resistance—the incomplete arrest of tumor cells by a drug. By interrogating the multidimensional protein state of individual cells, we demonstrate the phenomenon of fractional resistance both in a well-established ER+/HER2- model, T47D, as well as a primary tumor resected from a breast cancer patient. Through single-cell analysis, we found that tumor cells capable of proliferating in the presence of palbociclib showed unique combinations of enriched and depleted cell cycle regulators including cyclin B1, Cdt1, CDK2, and p21. Notably, resistant cells in both tumor models showed a common enrichment of the E2F1 transcription factor in G0 and G1 phases, suggesting that resistant cells may use a common molecular mechanism to overcome CDK4/6 inhibition. Tumor cells also showed an untimely enrichment of cyclin B in palbociclib treated cells, suggesting that this G2-associated cyclin might mediate drug escape. By tracing the trajectories of resistance in each tumor model, we visualize how plasticity in cell cycle regulators creates alternate cell cycle “paths” that allow some ER+/HER2- tumor cells to escape palbociclib treatment.

These findings explain several longstanding observations about ER+/HER2- breast cancer. For example, it has long been known that a correlation exists between the fraction of proliferating cells (i.e., Ki-67 staining) in a patient tumor and patient outcomes (23). Our work suggests that patients with more Ki-67-positive cells may have more plastic cell cycles that increase fractional resistance. Secondly, our work provides an alternative mechanism by which core cell cycle regulators can promote oncogenesis. Besides acquiring mutations in specific cell cycle genes, such as RB or cyclin E, we show that cell-to-cell variability in these same factors can promote drug resistance. Indeed, our observations of the enrichment of oncogenic protein factors may reflect the evolutionary pressure acting on tumor cells to select specific genetic alternations. Third, our work suggests a specific mechanism by which escaping tumor cells, through the accumulation of genetic events due to downregulation of the DNA licensing factor Cdt1, may be the seeds of genetically distinct tumor cells with more robust drug resistance. Supporting this idea, a recent study linked palbociclib-mediated arrest to downregulation of replisome components, defective origin licensing, and replication stress (73). Clinically, tumor mutational burden is associated with resistance to CDK4/6 inhibitors in patients with ER+/HER2- breast cancer (74) and deficient licensing due to downregulation of Cdt1 could be a key mechanism leading to accumulation of resistance promoting genetic events. Future work should investigate whether fractionally resistance cells are more prone to replication stress and genetic mutation.

Our work is consistent with the well-known additive effects of combining endocrine therapy and CDK4/6 inhibitors. If these drugs work together to reduce fractional resistance, then it may be profitable to consider additional combination therapies that further reduce the fractionally resistant subpopulation in ER+/HER2- breast tumors. Indeed, we found that the combination of palbociclib and tamoxifen, a drug that blocks estrogen receptor signaling, reduced fractional resistance in primary tumor cells (**Figure S6**). Finally, cells that are resistant to palbociclib showed higher levels of CDK2, implying that accumulating CDK2 may allow cells to overcome CDK4/6 mediated arrest. Therefore, CDK4/6 inhibitors with activity against CDK2 may have greater efficacy by also arresting this fractionally resistant subpopulation. In fact, patients with ER+/HER2- breast cancer treated with abemaciclib or ribociclib, which have greater activity against CDK2 compared to the more CDK4/6 specific palbociclib, have improved responses (75–78). Thus, using this human tumor 4i model we have uncovered a mechanism that explains clinical findings and supports the need for experimental systems using primary human tumors at the single-cell level to understand how tumors respond and resist therapies.

Evaluating therapeutic responses in terms of fractional resistance and cell cycle paths enhances our understanding of drug resistance mechanisms. Previous work has shown that cell cycle behaviors vary among tumors; here, we show that it also varies within the same tumor at a single-cell level. With the ability to fully profile the cell cycle behaviors in tumor cells—and distinguish among them—the field could make better predictions for when targeted therapy will work or how to develop new targeted treatments to arrest proliferating tumor cells. Future study will focus on understanding the range of sensitive and resistant cell cycle paths, and unique targetable drivers of resistant paths, both in ER+/HER2- tumors treated with CDK4/6 inhibitors, and in other human solid organ tumors. Identifying and characterizing tumor subpopulations with distinct sensitivities to targeted therapies could allow development of precision therapeutic regimens for individual patients based on specific tumor subpopulation drug sensitivities.

## MATERIALS AND METHODS

### Primary human breast tumor cells

Under an Institutional Review Board (IRB) approved protocol, we obtained a tumor sample from a female patient with invasive lobular carcinoma that was positive for expression of the estrogen receptor (ER+) and negative for amplification of HER2 (HER2-). The patient provided written informed consent. This tumor specimen was obtained in the operating room suite within 15 minutes of resection. The sample was placed in DMEM/F12 (Gibco) media with 1% Penicillin-Streptomycin and transferred immediately to the laboratory on ice. The tumor specimen was sharply minced into 2-4 mm fragments. Enzymatic dissociation was performed using Gentle Collagenase/Hyaluronidase (Stemcell Technologies Inc. 07919) in DMEM/F12 supplemented with 5% BSA, Hydrocortisone (Stemcell Technologies Inc.), HEPES (Corning), and Glutamax (Gibco) for 16 hours at 37°C with cell agitation. The cells were gently centrifuged and washed twice with PBS supplemented with FBS and HEPES buffer. Cells were resuspended in ammonium chloride solution (Stemcell Technologies Inc. 07800) and incubated at 37°C with 5% CO2 to remove red blood cells. Cells were centrifuged and briefly trypsinized in warm 0.05% Trypsin-EDTA (Gibco) and DNase I. Cells were centrifuged and washed then resuspended in DMEM/F12 with 10% FBS. Cells were then strained using multiple rounds of sequential straining with 100 um and 40 um cell strainers to remove cell debris. Cells were counted using fresh trypan blue and the Countess cell counter (Life Technologies). Cells were plated on a glass 96-well plate coated with poly-L lysine at 100,000 cells per well. Cells were allowed to adhere for 48 hours at 37°C with 5% CO2 in DMEM/F12 media with 10% FBS. After 24 hours, media and non-adherent cells were removed. DMEM/F12 media with 10% FBS was added containing vehicle, or palbociclib at 10 or 100 nM. Cells were incubated at 37°C with 5% CO2. After 24 hours of treatment, cells were fixed with PFA and 4i performed as described below.

FFPE slides sectioned at 4 microns were obtained from clinical pathology for the primary ER+/HER2- tumor. Immunohistochemistry (IHC) for Ki-67 antigen was performed using the Ki-67 Antibody (MIB-1, Dako) at 1:100 as we have previously described (79). A positive control was included. Ki-67 was scored according to the Ki-67 IHC MIB-1 pharmDx (Dako Omnis) Interpretation Manual for Breast Carcinoma. The Ki67 pharmDx score (%) was calculated as number of Ki-67 staining viable invasive (*in situ* disease was excluded) tumor cells divided by the total number of viable invasive tumor cells, multiplied by 100 for 2000 cells scored cells. The Ki67 staining for the primary tumor sample is shown in **Figure 1B**, which was scored as 14% Ki67+.

### Iterative immunofluorescence

We followed the protocol of Gut *et al.* (56) with the following modifications. T47D (ATCC HTB-133) or primary cells from human tumors were fixed by adding 8% PFA (Thermo Scientific cat#28908) directly to the samples (1:1 v/v with media) for a final concentration of 4% PFA and incubated for 30 minutes at room temperature (RT). Samples were rinsed 3 times with PBS (pH=7.4)(200 µL/well for 96 well format) and incubated with 0.1% Triton X-100 (50 µL/well)(Fisher cat#BP151) for 15 minutes at RT to permeabilize the cells for immunofluorescence. Samples were then rinsed a single time with PBS and then incubated with Hoechst (Sigma cat#94403)(50 µL/well; 1:2500 dilution in PBS) for 15 minutes at RT to stain the DNA contained in the nucleus of the cells. Cells were rinsed once with PBS,100 µL/well of PBS was added to the wells, and cells were imaged. This ‘pre-stain’ is a key first step as it ensures that 1) the cells are well distributed in the well and 2) serves as a necessary quality control step to ensure that the cells are suitable for 4i. Samples deemed suitable for 4i were eluted, even though labeling with a primary antibody has not occurred. This is done as the elution process further opens the cells and permits optimal labeling.

Elution of samples was carried out by first rinsing the samples three times with water. Elution buffer (EB) was prepared fresh from a pre-mix stock (L-Glycine [0.5M](Sigma cat#50046), Urea [3M](Sigma cat#U4883) and Guanidinum Chloride [3M](Invitrogen cat#15502-016)) combined with TCEP-HCl [70 mM](Sigma #646547) and HCl (Fisher cat#SA49) to obtain a pH to 2.5. Samples were washed three times with EB (50 µL/well) for 10 minutes at RT with gentle shaking. Of note, it is important not to exceed the number of washes or the duration of the washes as this may degrade the samples. Once elution was complete, the sample was rinsed one time with PBS prior to labeling with primary antibodies.

Labeling with primary antibodies first requires incubation with sBS (4i blocking solution) for 1 hour at room temperature (50 µL/well). The blocking solution was made up fresh and for every mL of solution one adds 14.6 mg Malemide [100 mM](Sigma cat#129585) and 5.35 mg NH_4_Cl [100 mM](Sigma cat#A9434) to conventional blocking solution (cBS)(1% BSA (Sigma cat#A7906) in PBS). Once incubation with blocking solution was complete, samples were rinsed one time with PBS and primary antibodies (50 µL/well) were applied for an overnight incubation at 4°C with gentle rocking/shaking. It is important to note that the antibody solution was made in a conventional blocking solution at a dilution that is empirically determined and may contain several different antibodies. This does not present an issue as long as the antibodies have different species of origins (see **Table S1** for a list of primary antibodies used in this study). Alternatively, samples may be incubated with the primary antibody solution at room temperature for an hour or more, but labeling may not be as robust. Once the incubation with the primary antibody solution was complete, samples were rinsed one time with PBS, followed by three washes with PBS for 5 minutes each, followed by one final rinse in PBS. The final rinse with PBS was carefully aspirated off the sample to ensure all residual antibodies had been removed. Immediately following incubation with primary antibodies, fluorescent secondary antibodies specifically directed at the primary antibodies were applied. We used the Alexa series of secondary antibodies at a dilution of 1:500 in cBS along with Hoechst DNA stain at 1:2500. The secondary solutions (50 µL/well) were incubated for 1 hour at RT with gentle rocking and under conditions excluding light to prevent any photobleaching of the secondary fluorophores. Once this step was complete, cells were rinsed/washed in the same exact manner as the end of the primary antibody incubation step. During the wash step, fresh imaging buffer (IB) was prepared, which consists of N-acetylcysteine (NAC, Sigma cat#A7250) in water at a final concentration of 700 mM and pH of 7.4. We added 100 µL/well of IB to the samples and immediately imaged the cells.

Imaging was performed on a Nikon TiE inverted microscope utilizing a plan apo lambda 20X objective lens (NA = 0.75) with an Andor Zyla 4.2P sCMOS camera as a detector. NIS-Elements HCA (high content analysis) JOBS software was utilized in the acquisition of images as it permits the imaging of entire wells in a fast and automated fashion. Upon completion of imaging, samples were eluted per the protocol described above and the next round of labeling and imaging was performed. It should be noted that every other round after elution, and before the next round of labeling, samples were imaged with the same exact experimental parameters with successful elution. This results in little to no fluorescent signal and ensures that the antibodies from the previous round have been successfully removed via the elution process and that no residual labeling is present to ‘contaminate’ the next round of imaging. This process of imaging and elution was repeated in an iterative manner to build a molecular profile for individual cells for each sample and treatment condition.

### Image processing and cell property quantification

The image processing pipeline consisted of several steps to convert the raw images in the Elements nd2 format to a matrix of single cells with protein expression quantified in the nucleus, ring, and cytoplasm. The four primary steps were: 1) **cell segmentation** via the Cellpose algorithm (71) to define the nucleus for each round from the Hoechst staining, 2) cell segmented masks were **aligned** across all rounds of images, 3) **punchmasks** were manually drawn to exclude any debris (cellular or otherwise), and 4) **cell properties** were calculated from individual segmented nuclei for all the intensity channels. We followed the image preprocessing pipeline as described in the GitHub repository: https://github.com/fjorka/4i_analysis.

### Data preprocessing

To compare tumor cells from a cell culture model of ER+ breast cancer (T47D) and tumor cells from a primary tumor sample resected from a patient, we performed a series of preprocessing steps. Following image preprocessing and cell property quantification, we computationally filtered cells within the primary tumor sample to retain only the tumor epithelial cells by gating cells according to the median expression of ER and PR (see **Figure S4**). Feature selection was then performed by selecting the intersection of core cell cycle regulators profiled in both datasets (*P =* 14). Lastly, T47D and primary tumor datasets were standardized independently by mean centering and scaling to unit variance. The abbreviated experimental pipelines for the T47D and primary tumor samples are shown in **Figure 2A** and **Figure S3**, respectively.

### Cell cycle annotations

The bimodal distribution of the ratio of phosphorylated to total RB levels (pRB/RB) was used to distinguish proliferative cells (G1/S/G2/M, high pRB/RB) from arrested cells (G0, low pRB/RB). To agnostically set the pRB/RB threshold for both datasets without any underlying assumptions on the shape or spread of the distribution, we implemented a data normalization step outlined previously in Ref. (80), based on the idea that if a distribution is bimodal, there will be a region of higher density on one side of the median as compared to the other. More precisely, given a sorted list of expression values, *x_pRB_*_/*RB*_, we first computed the median of the distribution as *m =* median(*x_pRB_*_/*RB)*_). We then folded the left side of the distribution, *x_pRB_*_/*RB*_ < *m,* over the right side of the median by *z_pRB_*_/*RB*_ [*x_pRB_*_/*RB*_ < *m*] *= 2*m* – *x_pRB_*_/*RB*_ [*x_pRB_*_/*RB*_ *< m*], where *z* is the new one-sided distribution. Next, we computed a specified percentile, *p,* of this one-sided distribution and subtracted the median, denoting this difference as *a*, *z_(pRB_*_/*RB,*_ *_p)_* – *m* = *a*. The cutoff point of the second mode of the distribution (i.e., proliferative cells with high pRB/RB) was then defined according to the values of *x_pRB_*_/*RB*_ that fell within the range (*m – a*, *m + a*). More specifically, we denote *s* = |(*z* ∈ (*m – a*, *m + a*))|, where | is the cardinality of the set within the specified range. We define the point separating the modes of the distribution, *c,* as *c* = *s / n*, where *n* is the total number of cells in the distribution. We selected a percentile value of (*p* = 0.2) for T47D, (*p* = 0.7) for T47D replicate, and (*p* = 0.7) for the primary tumor sample based on the distribution of pRB/RB expression values (See **Figures 2B, S1A, S5**).

For the proliferative cells, as indicated above by high expression of pRB/RB, cell cycle phase annotations (G1, S, and G2/M) were subsequently determined by fitting a three component Gaussian Mixture Model to the log-transformed measurements of DNA content, cyclin A, and cyclin B1. Unsupervised clusters were annotated as follows: G1 (DNA content = 2C, low cyclin A), S (DNA content = 2-4C, medium cyclin A), and G2/M (DNA content = 4C, high cyclin A). The Gaussian Mixture Model was implemented using the sklearn 0.24.1 package in Python.

### Sketching

To identify a limited subset of representative cells for each dataset and facilitate a direct comparison cell cycle states across samples and treatment conditions, we selected an equal number of cells (*n* = 2,000) from each treatment condition (untreated, 10 nM, and 100 nM palbociclib) within a dataset (T47D, primary tumor) using kernel herding sketching (57). Kernel herding sketching performs principled downsampling of the data and selects prototypical cells that are representative of the original distribution of cell type frequencies (e.g., cell cycle phases), while also ensuring rare cell types are sufficiently sampled. For each dataset, sketched cells from each condition were then vertically concatenated into a *N* × *P* matrix prior to downstream analysis, where *N* is the number of sketched cells across three treatment conditions (*N =* 6,000) and *P* is the number of profiled proteomic imaging features (*P =* 14).

### Confidence intervals

To identify shared and distinct mechanisms of resistance to palbociclib treatment, we examined the fractional arrest profiles of proliferating T47D and primary tumor cells as follows. For each dataset (T47D, primary tumor), we computed two sample *t-*tests assuming equal variance between each cell cycle effector in untreated proliferative cells and each treatment condition separately. More specifically, we computed 95% confidence intervals between proliferative untreated and 10 nM palbociclib cells, and 95% confidence intervals between proliferative untreated and 100 nM palbociclib cells.

### Logistic regression

Logistic regression (81) is a supervised learning algorithm that can be used to predict the probability of a binary outcome (e.g., control, treated) based on a set of input features (e.g., proteomic imaging features). To ascertain changes in cell cycle regulators associated with palbociclib treatment, a logistic regression model was trained on the proteomic expression profiles of cells within each cell cycle phase (G0, G1, S, G2/M) for each dataset (T47D, primary tumor) to predict the treatment group of a tumor cell (control, treated). In this case, the control group consisted of untreated cells, whereas the treated group consisted of cells treated with either concentration of palbociclib (10 nM, 100 nM). For each dataset and phase, nested ten-fold cross-validation was performed using stratified random sampling to assign cells within a particular phase to either a training or a test set. Using a grid search, hyperparameters were tuned within each fold prior to training the model, and cells were classified as control or treated from the test data. Classification performance was subsequently assessed by computing the area under the receiver operator characteristic curve (AUC ROC). Logistic regression was implemented using the sklearn v0.24.1 package in Python.

### Data integration with TRANSACT

TRANSACT (Tumor Response Assessment by Nonlinear Subspace Alignment of Cell lines and Tumors) (69) is a nonlinear data integration method that can be used to identify a shared subspace of preclinical cell lines and patient-derived samples. Briefly, TRANSACT merges datasets by performing kernel principal components analysis (82) on each individual dataset, and then geometrically aligns these nonlinear principal components to extract principal vectors that represent similar nonlinear weighted combinations of expression profiles across data samples. A consensus data representation, corresponding to biological processes that are present within both preclinical cell lines and primary tumor samples, is then computed by optimizing the match between interpolated sets of principal vectors using geodesic flow (83). We performed data integration of T47D and primary tumor samples using TRANSACT to more robustly represent and compare cell cycle trajectories under palbociclib treatment. More specifically, we identified a shared latent space by first computing consensus features for T47D and primary tumor samples, and then projecting both datasets onto the consensus features. Here, the integrated dataset, *F^T^*^47*D,Tum*^, consisted of 12,000 cells and 14 shared consensus features. Of note, integration was performed on the sketched datasets to ensure that the joint latent space was not overwhelmed by one data modality when performing downstream analyses, such as dimensionality reduction and trajectory inference. TRANSACT was implemented using the transact-dr v1.0.1 package in Python, where cell similarity was defined using a radial basis function with a scaling factor, 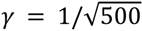.

### PHATE dimensionality reduction

To visualize high dimensional single-cell 4i profiles of the cell cycle, we performed nonlinear dimensionality reduction with PHATE (Potential of Heat-diffusion for Affinity-based Trajectory Embedding) on the integrated dataset of T47D and primary tumor samples. PHATE (70) is a nonlinear dimensionality reduction method that effectively represents the geometry of complex continuous data structures and has been shown previously (52, 53, 84) to successfully recapitulate proliferative and arrest cell cycle trajectories. PHATE was implemented using the phate v1.0.7 package in Python by constructing a *k*-nearest neighbor graph (*k =* 150) according to pairwise Euclidean distances between all pairs of cells from the consensus feature space computed by TRANSACT, *F^T^*^47*D,Tum*^.

### Trajectory inference and alignment

To characterize trajectories through the cell cycle under palbociclib treatment, we performed trajectory inference using Slingshot (71) on each dataset (T47D, primary tumor) and treatment condition (untreated, 10 nM, and 100 nM palbociclib). This trajectory inference method was chosen as it was shown previously (85) to outperform alternative methods on inferring simple continuous or branched cellular trajectories. Slingshot was implemented using the slingshot v2.7.0 package in R by 1) fitting a minimum spanning tree through cluster centroids defined by cell cycle phase annotations, and then 2) estimating pseudotime by projecting cells onto the principal curves fit through the PHATE embedding generated from the consensus feature space computed by TRANSACT. The *root* (starting) cluster was defined as the G0 phase. Across most inferred cellular trajectories, Slingshot identified the canonical ordering of cell cycle phases (G0 to G1 to S to G2/M). However, we note that in two scenarios (untreated T47D and 100 nM palbociclib primary tumor), Slingshot identified a minimum spanning tree spanning from G0 to G1 to G2/M phases for the primary tumor and G0 to S to G2/M for the T47D, respectively.

Given that trajectory inference was performed on cells from each treatment condition separately, we subsequently aligned the trajectories onto one common pseudotime axis using TrAGEDy to enable a direct comparison of continuous proteomic expression profiles across treatment conditions. TrAGEDy (Trajectory Alignment of Gene Expression Dynamics) (72) is a trajectory alignment method that can align cells from two independently generated trajectories and has been shown previously to enable robust comparisons of continuous expression trends across treatment conditions when aligning Slingshot trajectories from PHATE dimensionality reduced single-cell data. Methodologically, TrAGEDy first interpolates points at different regions of the trajectory to overcome any noise inherent to single-cell data. Next, the Spearman correlation is computed between the set of interpolated points along the two trajectories to define a trajectory similarity matrix. Lastly, TrAGEDy uses a dynamic time warping approach (86) with modifications to account differences in cell states in order to find the optimal alignment through the similarity matrix of interpolated points. This approach ensures that the original pseudotemporal ordering is preserved, while the distance between points across trajectories is minimized. For each dataset, we performed trajectory alignment with TrAGEDy by aligning the 10 nM and 100 nM trajectories to one another, followed by alignment to the untreated trajectory. TrAGEDy was implemented with 50 interpolating points using the R code provided in the GitHub repository at: https://github.com/No2Ross/TrAGEDy.

To visualize continuous feature expression trends, a generalized additive model (GAM) with a cubic spline basis function with shrinkage was fit for each feature as an outcome along the aligned pseudotime as sole covariate using the mgcv v1.8-42 package in R. Moreover, to identify an approximate transition point from arrest into proliferation, we computed the inflection point where approximately 50% of the cells were G0 and 50% of the cells were proliferative (non-G0) for each trajectory. To do so, we discretized the aligned pseudotime values into bins and then computed the ratio of G0/non-G0 cells for each bin. The transition point was defined as the aligned pseudotime value where this ratio was approximately one. For the untreated trajectories, we chose a smaller number of bins (*n =* 25) to find the inflection point due to the larger number of proliferative cells, whereas for the treated trajectories, we chose a larger number of bins (*n =* 50). Of note, this transition point was excluded for the 100 nM palbociclib primary tumor trajectory due to the small sample size of proliferative cells.

## DATA AND CODE AVAILABILITY

Preprocessed single-cell 4i datasets are publicly available in the Zenodo repository: https://doi.org/10.5281/zenodo.7930054. Source code for image preprocessing, including cell segmentation, transformation, alignment, and quantification are publicly available in the GitHub repository: https://github.com/fjorka/4i_analysis. Source code for computational analyses, including functions for preprocessing, sketching, integration, trajectory inference, and other computational analyses as described in this manuscript are publicly available in the GitHub repository: https://github.com/purvislab/fractional_resistance.

## ACKNOWLEDGMENTS

We thank the patient who donated tissue for this study. We thank Kasia Kedziora for her expertise in establishing the imaging processing pipeline. Lastly, we thank Wayne Stallaert and Jean Cook for their insightful discussions relating to the cell cycle. This work was supported by NIH F31HL156464 (T.M.Z.), NIH F31-HL156433 (J.S.R.), NIH 5T32-GM067553 (J.S.R.), NIH P50CA058223 (P.M.S.), R01-GM138834 (J.E.P.), NSF CAREER Award 1845796 (J.E.P.), and NSF Award 2242980 (J.E.P.). This work was also supported in part by P30CA016086 UNC Lineberger Comprehensive Cancer Center Core Support Grant and funding from the UNC School of Medicine for the Computational Medicine Pilot Award Program (P.M.S., S.C.W.).

## AUTHOR CONTRIBUTIONS

T.M.Z, S.C.W., P.M.S., and J.E.P. conceptualized the study. P.M.S. performed tumor resection. A.A.W. and S.C.W. processed primary tumor cells. S.C.W performed cell culture, 4i protocol and image preprocessing with help from A.N. and H.D. T.M.Z. and J.S.R. performed all computational analyses. M.R.K. advised on statistical analyses. Writing and manuscript preparation was primarily handled by T.M.Z., P.M.S., and J.E.P. All authors read and approved the final manuscript.

## DECLARATIONS OF INTEREST

The authors have no competing declarations of interest.

## Notes

### Competing Interest Statement

The authors have declared no competing interest.

