## Supplementary Information for "Cell cycle plasticity underlies fractional resistance to palbociclib in ER+/HER2- breast tumor cells"

| Primary Antibody | Species | Vendor/Cat# | Dilution |
|---|---|---|---|
| CDK2 | Goat | R&D Systems/AF4654 | 1:100 |
| CDK4 | Rabbit | Abcam/ ab108355 | 1:400 |
| CDK6 | Rabbit | Abcam/ab124821 | 1:250 |
| Cdt1 | Rabbit | Cell Signaling Technology (CST)/8064 | 1:200 |
| Cyclin A | Mouse | Santa Cruz/sc-271682 | 1:50 |
| Cyclin B1 | Goat | R&D Systems/AF6000 | 1:100 |
| Cyclin D1 | Mouse | Santa Cruz/sc-20044 | 1:100 |
| Cyclin E | Mouse | Santa Cruz/sc-247 | 1:50 |
| E2F1 | Mouse | Santa Cruz/sc-251 | 1:100 |
| Estrogen Receptor (ER) | Rabbit | Abcam/ab32063 | 1:200 |
| Ki67 | Rabbit | Abcam/ab15580 | 1:800 |
| p16 | Rabbit | Abcam/ab108349 | 1:400 |
| p19 (CDKN2D) | Rabbit | Invitrogen/PA5-83665 | 1:200 |
| p21 | Goat | R&D Systems/AF1047 | 1:200 |
| PanCK | Mouse | CST/4545 | 1:200 |
| Phospho-Retinoblastoma (pRB) | Rabbit | CST/8516 | 1:1000 |
| Progesterone Receptor (PR) | Mouse | Thermo Fisher/MA5-12658 | 1:400 |
| Retinoblastoma (RB) | Mouse | CST/9309 | 1:500 |

**Table S1**. List of primary antibodies utilized in the 4i rounds of imaging along with host species, vendor information and dilution utilized in the experiment.

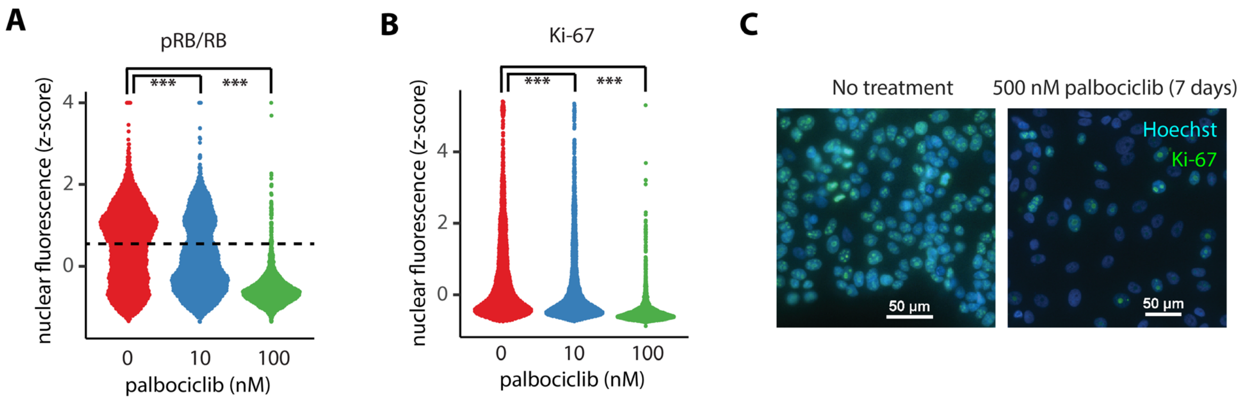


**Figure S1. Fractional resistance in a biological replicate and under long-term treatment with palbociclib. A.** Distribution of pRB/RB in T47D cells under 0, 10, or 100 nM palbociclib. The dotted line marks the statistically determined threshold for demarcating cells as proliferating despite palbociclib treatment. Cells above the dotted line were used for characterizing proliferative cells. **B.** Distribution of nuclear Ki-67 levels at 0, 10, or 100 nM palbociclib. *** indicates a *P*-value < 0.001 using a two-sided Kolmogorov-Smirnov test between untreated and treated cells. **C.** The ER+/HER2- cell line MCF7 was treated with 500 nM palbociclib for 7 days. Despite a reduction in growth rate, a significant fraction of cells (~12%) retained pRB/RB and KI-67 expression.

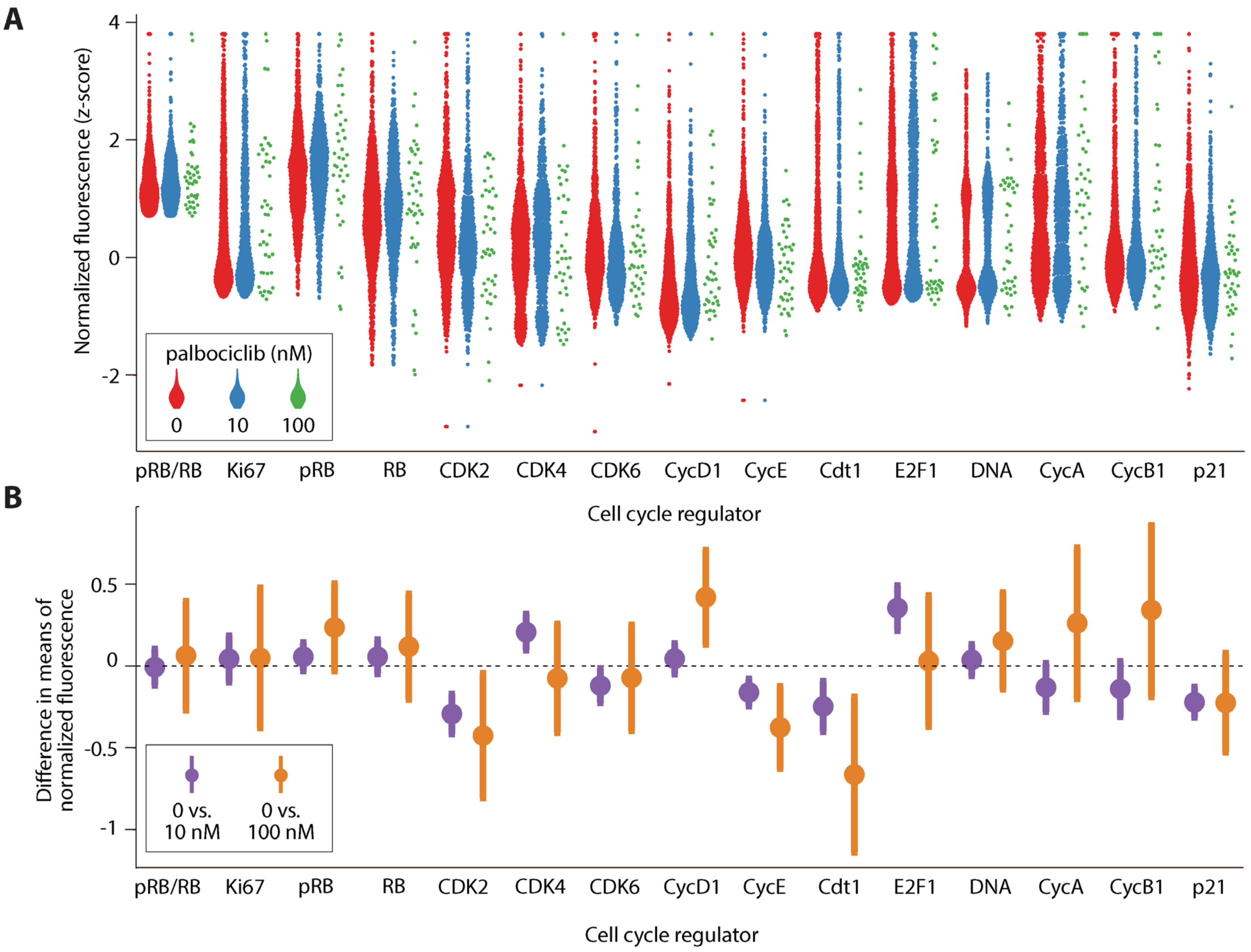


(Figure S2 continued on next page)

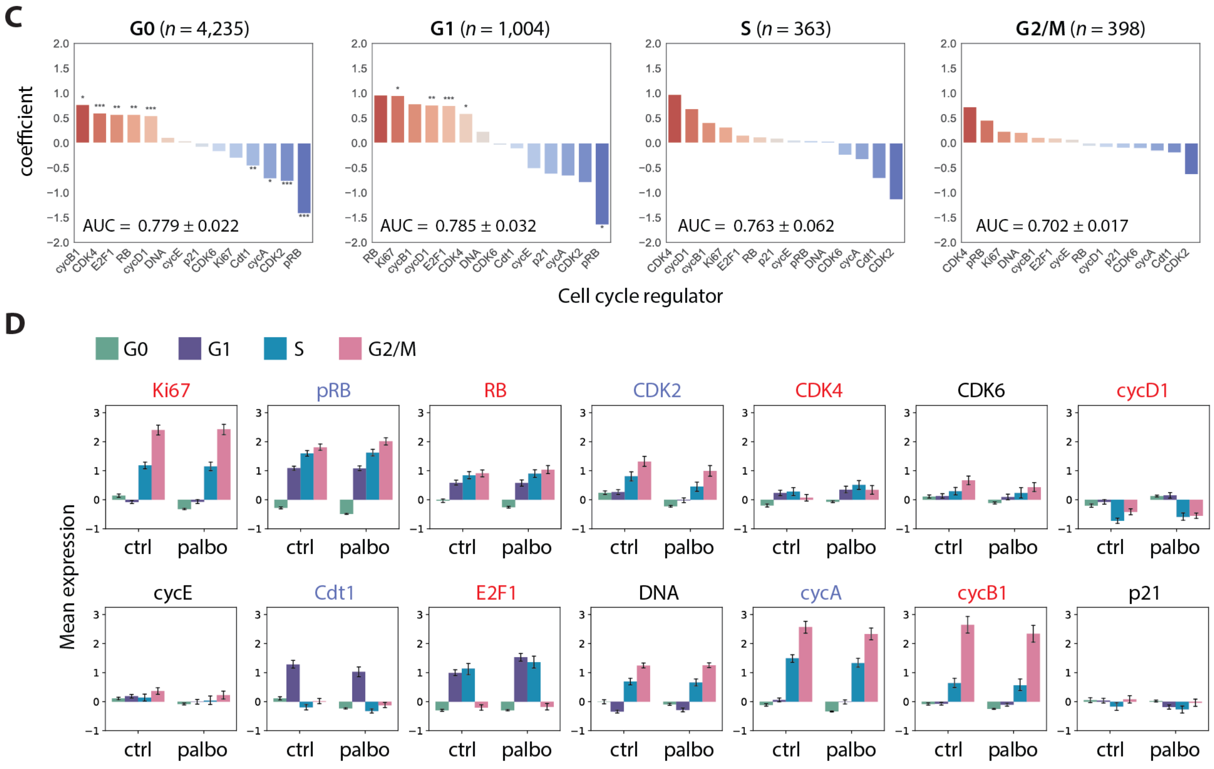


**Figure S2. Shifts in expression of cell cycle proteins among fractionally resistant tumor cells for T47D biological replicate. A.** Beginning with a downsampled dataset containing 2,000 cells from each condition (see **MATERIALS AND METHODS**), we identified the proliferating cells using the pRB/RB threshold defined in **Figure S1A**. Proliferating single-cell distributions of cell cycle regulators at 0 nM, 10 nM, or 100 nM palbociclib, showing an expected reduction in the number of cells at higher drug concentrations. **B.** 95% confidence intervals (CI) of proliferating cells for the differences in mean expression (normalized z-scores) between either untreated cells and 10 nM palbociclib (*purple*); or untreated and 100 nM palbociclib (*orange*). Confidence intervals overlapping with the dashed line at 0 indicate a lack of statistical significance. Confidence intervals are wider when comparing 0 vs. 100 nM due to lower sample sizes at the highest dose of palbociclib. **C.** Logistic regression on all 6,000 cells predicting the odds that a given cell is either untreated (0 nM) or treated (10 nM or 100 nM) based on expression of its cell cycle regulators. Cell-to-cell increases in regulators shown in red (e.g., CDK4), or decreases in regulators shown in blue (e.g., CDK2), increase the odds of association with treated (10 nM or 100 nM) versus untreated cells. A separate regression was performed for each phase (G0, G1, S, G2/M), where the last three are considered proliferating (high pRB/RB). This analysis was performed on all cells, including non-proliferating (G0) cells. Significance: *, $P < 0.05$; **, $P < 0.01$, ***, $P < 0.001$. **D.** Expression levels of cell cycle features, stratified by cell cycle phase for all 6,000 cells. Bar height is the mean expression for untreated (ctrl) or palbociclib-treated (10 nM and 100 nM) cells. Error bars represent confidence intervals.

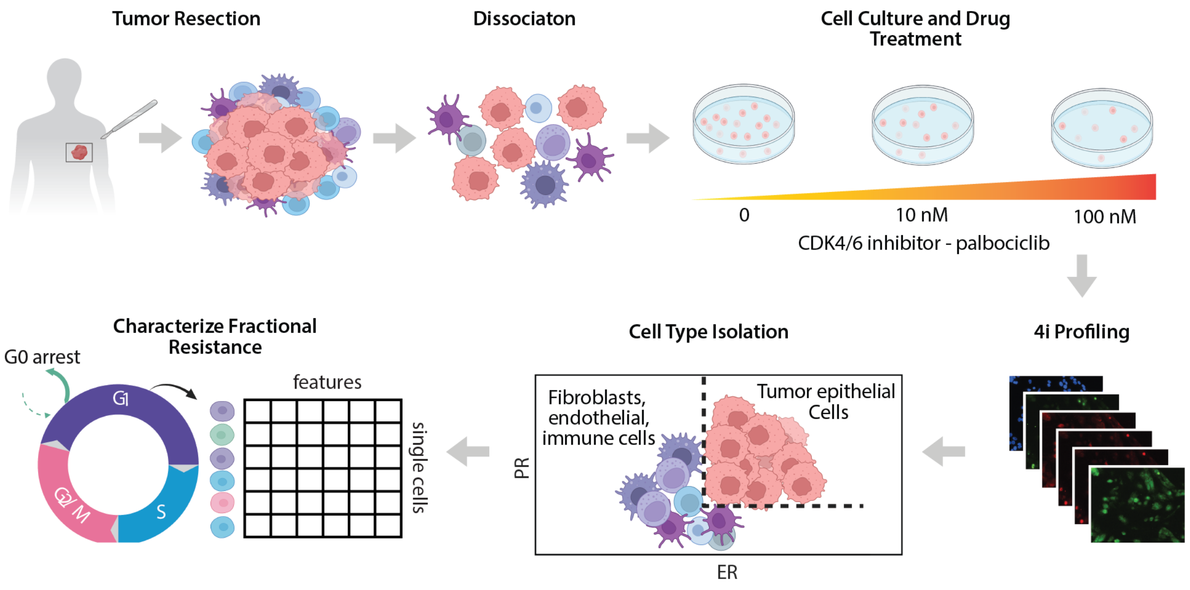


**Figure S3. Experimental and analysis pipeline for characterizing fractional resistance in primary tumor cells.** Detailed methods may be found in the **MATERIALS AND METHODS** section of the main text.

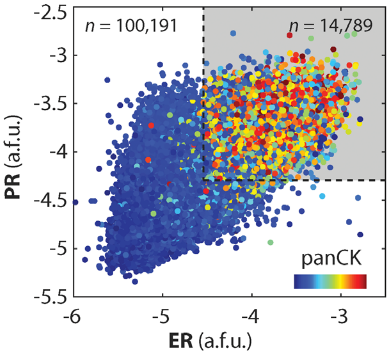


**Figure S4. Image-based isolation of ER+/PR+ cells from resected tumor specimen.** ~15% of cells were identified as ER+ and PR+ and subjected to downstream analysis.

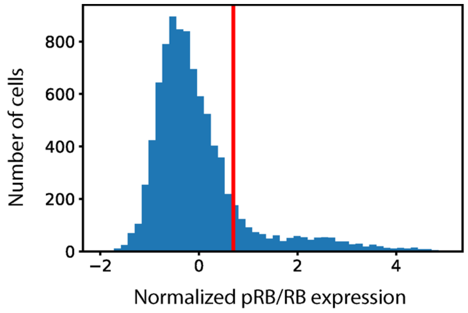


**Figure S5. Distribution of pRB/RB in primary ER+/HER2- tumor cells.** The red line marks the statistically determined threshold for demarcating cells as proliferating despite palbociclib treatment. Cells to the right of the line were used for downstream analyses.

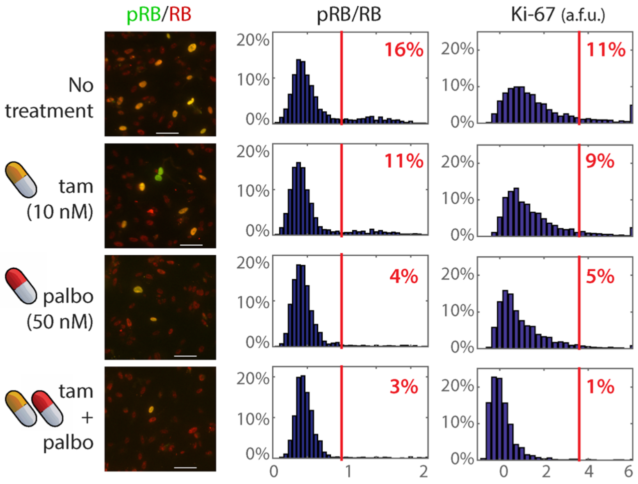


**Figure S6. Fractional resistance in resected tumor cells under combination therapy.** Primary ER+/PR+ tumor cells were treated with 10 μM tamoxifen and/or 50 nM palbociclib prior to 4i profiling. Y-axis denotes the percentage of cells at a given expression level.
